# Differential Cytotoxicity of PVP-Copper Nanoparticles in Breast Cancer Cell Lines: Insights from Basal Transcriptomic Profiles

**DOI:** 10.64898/2026.09.21.753010

**Authors:** Anelise G. Silva, Izaque J. O. Silva, Pedro P. Tanaka, Matheus Z. Monteiro, Sérgio Servilha Oliveira Filho, Jefferson M. Souza, Valeria S. Marangoni, Sandra M. G. Dias, Cyro. Z. V. Negrão

## Abstract

Breast cancer is a heterogeneous disease comprising molecular subtypes with distinct therapeutic vulnerabilities. Among emerging therapeutic strategies, copper-based nanoparticles have shown anticancer activity. However, whether these responses differ across breast cancer subtypes and which molecular programs underlie copper sensitivity remain poorly understood. We investigated subtype-specific responses to Polyvinylpyrrolidone-assisted copper nanoparticles (CuNP-PVP) in breast cancer cell lines and the molecular programs underlying copper adaptation. CuNP-PVP were synthesized and evaluated in luminal (MCF-7), HER2-positive (SKBR3), and triple-negative (MDA-MB-231) breast cancer cell lines, with transcriptomic analyses integrated with CCLE and TCGA-BRCA datasets. UV-Vis spectroscopy revealed a plasmon resonance band at 591 nm, while TEM showed spherical nanoparticles with an average diameter of 76.1 ± 21.0 nm. DLS indicated a larger hydrodynamic diameter, consistent with PVP coating; ζ-potential measurements showed a surface charge of −13 mV, and XPS confirmed mixed reduced and Cu(I)/Cu(II) oxidation states. Exposure to 100 μg/mL CuNP-PVP induced subtype-dependent effects on cell viability, with MCF-7 showing greater sensitivity, whereas MDA-MB-231 and SKBR3 were less affected. Baseline transcriptomic analyses revealed enrichment of oxidative stress, metabolic adaptation, and lysosomal programs in less sensitive cells, whereas MCF-7 showed higher expression of lipoylation-related genes. These features supported the development of a Potential Biological Copper-Response Index (PBCRI), which captured subtype-associated transcriptional patterns across independent cell lines and patient tumors, with luminal models showing lower scores and basal/TNBC models showing higher scores. Together, these findings identify candidate transcriptional programs associated with subtype-specific responses to CuNP and provide a framework for investigating potential molecular determinants of copper sensitivity.

**Highlights:**

- PVP-assisted copper nanoparticles showed mixed metallic/oxidized Cu phases.
- CuNP-PVP induced cell-dependent cytotoxicity in breast cancer cells.
- MCF-7 cells were more sensitive to CuNP-PVP than TNBC and HER2+ cells.
- Baseline profiles differed in lipoylation, antioxidant, and lysosomal programs.
- PBCRI captured subtype-associated copper-response transcriptional patterns.

## Introduction

Breast cancer remains one of the leading causes of cancer-related mortality among women worldwide [1]. It is a highly heterogeneous disease characterized by subtype-specific differences in gene expression, metabolic activity, and therapeutic response [2,3]. Clinically, breast tumors are commonly classified into luminal A, luminal B, HER2-enriched, and triple-negative (TNBC) subtypes [4]. Each subtype exhibits distinct molecular signatures and biological behaviors. They also exhibit distinct patterns of mitochondrial activity [5]. These differences are accompanied by variations in redox regulation, mitochondrial metabolism, and cellular stress-response pathways, which may influence how tumor cells adapt to metabolic perturbations [6,7].

Recent studies have identified transition metal homeostasis, including copper metabolism, as a regulator of tumor growth and cellular responses [8,9]. Copper is an essential trace metal involved in mitochondrial respiration, antioxidant defense and multiple enzymatic processes through its incorporation into cuproproteins [10]. Because of its intrinsic redox activity, cells must tightly regulate copper levels through coordinated transport and trafficking systems [11]. In cancer cells, dysregulated copper homeostasis has been linked to enhanced proliferative signaling, angiogenesis and metastatic potential [12]. However, excessive intracellular copper accumulation can become highly cytotoxic, requiring cellular mechanisms that maintain copper homeostasis and tolerance [8].

Copper-induced cytotoxicity involves multiple interconnected biological processes, including oxidative stress, mitochondrial dysfunction, proteotoxic stress, and alterations in lysosomal and autophagic pathways [13]. The relative contribution of these mechanisms depends on cellular metabolic state and antioxidant capacity, which influence the ability to adapt to copper overload [14]. Among the mechanisms proposed, cuproptosis, first described by Tsvetkov et al. in 2022 [15] is a regulated form of copper-dependent cell death in which copper binds lipoylated proteins of the tricarboxylic acid (TCA) cycle, promoting protein aggregation and mitochondrial dysfunction [16–18]. However, accumulating evidence suggests that copper toxicity cannot be explained solely by cuproptosis and instead reflects the coordinated activation of multiple stress-response pathways [19]. Because these responses are closely linked to cellular metabolism, metabolic heterogeneity across breast cancer subtypes may contribute to differential sensitivity to copper [20]. Luminal tumors generally exhibit greater dependence on oxidative phosphorylation, whereas TNBCs frequently display enhanced metabolic plasticity and glycolytic adaptation [21,22].

To exploit the cytotoxic potential of copper, considerable effort has been devoted to developing systems capable of increasing intracellular copper levels in cancer cells [23–25]. These strategies include nanotechnology-based delivery systems that enable controlled copper transport and enhanced intracellular accumulation [26,27]. Copper-based nanomaterials have been widely investigated in cancer models, including breast cancer cell lines, as part of efforts to explore their cytotoxic effects and therapeutic potential [28]. Within this context, copper nanoparticles (CuNPs) have emerged as promising candidates, as their redox activity enables efficient intracellular copper delivery and sustained cytotoxic responses [29–31]. However, their biological effects are strongly influenced by physicochemical characteristics, including particle size, morphology, oxidation state, and colloidal stability [32]. Because CuNPs are highly susceptible to oxidation, surface stabilization is essential for preserving their physicochemical characteristics and biological activity [33,34]. Polyvinylpyrrolidone (PVP) is widely used as a stabilizing agent because it reduces aggregation and oxidation while improving colloidal stability in biological media [35,36].

Despite recent advances in copper-based nanotherapeutics for breast cancer, molecular determinants of sensitivity to copper nanoparticle exposure remain poorly defined. Most studies have investigated copper or copper oxide nanoparticles using one or a limited number of breast cancer cell lines [37–39], limiting the ability to capture biologically relevant differences across molecular subtypes. As a result, current models do not resolve how subtype-specific differences in metabolism, mitochondrial function, and copper handling shape cellular responses to CuNPs. Consequently, the contribution of nanoparticle properties versus intrinsic cellular programs to copper-induced cytotoxic responses remains unclear [40].

In this paper, we investigated the cytotoxic effects of PVP-assisted copper nanoparticles (CuNP-PVP) across distinct breast cancer cell lines. By combining comprehensive physicochemical characterization with biological evaluation, we assessed subtype-specific differences in susceptibility to CuNP-PVP exposure. We further integrated transcriptomic analyses from experimental models, CCLE, and TCGA-BRCA datasets to identify molecular programs associated with copper adaptation and to determine whether these features could explain subtype-specific responses to copper-induced stress.

## 2. Experimental Section

### 2.1. Materials

Copper (II) chloride dihydrate (CuCl₂·2H₂O, MW = 170.48 g·mol⁻¹), ascorbic acid (C_6_H_8_O_6_, MW = 176.12 g·mol⁻¹), sodium borohydride (NaBH₄, 96%, MW = 37.83 g·mol⁻¹), acetone, RPMI-1640 medium, and Dulbecco’s Modified Eagle Medium (DMEM) were purchased from Sigma-Aldrich. Polyvinylpyrrolidone (PVP K-30, MW ≈ 40,000 g·mol⁻¹) was obtained from Dinâmica. Absolute ethanol was purchased from Merck KGaA, and isopropyl alcohol was obtained from Synth. Fetal bovine serum (FBS) was purchased from Vitrocell, and penicillin–streptomycin solution, as well as the CyQUANT™ MTT Cell Proliferation Assay Kit, were obtained from Thermo Fisher Scientific. Ultra-low attachment plates were purchased from Corning Costar. Deionized water was used in all preparations, and phosphate-buffered saline (PBS) was prepared in-house. All reagents were used as received without further purification.

### 2.2 Synthesis of Copper Nanoparticles

CuNPs were synthesized using a modified procedure based on previously reported studies by Sharma et al. [41]. Briefly, PVP (5 mM) was dissolved in 17.5 mL of deionized water at 65 °C under constant magnetic stirring. CuCl₂ (11.6 mM, 2.5 mL) was subsequently added, and the reaction temperature was increased to 80 °C. An ice-cooled NaBH₄ solution (100 mM, 2.5 mL) was added dropwise under continuous stirring. The reaction mixture was immediately transferred to an ice bath and maintained for 21 min. The suspension was then reheated to 72 °C for 15 min, followed by the slow addition of ascorbic acid (200 mM, 2.5 mL). Nanoparticle formation was monitored by the color transition from yellow-green to light orange, which occurred after approximately 4–5 min. The reaction mixture was maintained at room temperature for 1 h prior to purification. For purification, 1 mL aliquots of the suspension were centrifuged at 20,000 rcf for 20 min at 4 °C. The supernatant was discarded, and the pellet was washed with absolute ethanol followed by an additional centrifugation step at 20,000 rcf for 30 min at 4 °C. The final pellet was redispersed in autoclaved deionized water to obtain purified CuNP-PVP.

### 2.3 Characterization Techniques

UV–Vis spectra were acquired at room temperature using a UV-2600/2700i spectrophotometer (Shimadzu) equipped with an integrating sphere accessory. Measurements were performed in quartz cuvettes using aqueous dispersions diluted 1:1 in deionized water. Dynamic Light Scattering (DLS) and ζ-potential measurements were conducted using a Zetasizer Nano ZS90 µV (Malvern Instruments) at 25 °C. Prior to analysis, samples were diluted 1:10 in ultrapure water. Approximately 30 µL of each sample was loaded into quartz cuvettes for hydrodynamic size measurements, whereas ζ-potential was measured in capillary cells (∼1.5 mL).

Fourier Transform Infrared Spectroscopy (FTIR) measurements were performed using an IRSpirit-T spectrophotometer (Shimadzu) equipped with a QATR-S attenuated total reflectance accessory. For sample preparation, 8 mL of the CuNP-PVP was concentrated to a final volume of 1 mL in water and dried under vacuum using a SpeedVac MIVAC DUO system (Remma). Spectra were processed using Python-based routines for Savitzky–Golay smoothing, normalization, and baseline correction. X-ray Photoelectron Spectroscopy (XPS) analyses were carried out using a K-Alpha spectrometer (Thermo Scientific). Approximately 30 µL of purified CuNP-PVP suspension was deposited onto silicon substrates and dried at room temperature prior to analysis. The resulting spectra were processed using CasaXPS (Version 2.3.27). Background correction was performed using a Shirley function, followed by peak deconvolution using a Gaussian–Lorentzian (GL) line shape.

Scanning Electron Microscopy (SEM) analyses were performed using purified CuNP-PVPs deposited onto conductive Si substrates and dried at room temperature prior to imaging. Transmission Electron Microscopy (TEM), High-Resolution Transmission Electron Microscopy (HRTEM), and Scanning Transmission Electron Microscopy (STEM) analyses were performed using a JEOL JEM-2100F microscope operated at an accelerating voltage of 200 kV. CuNP-PVP were deposited by drop-casting 5 µL of dispersions prepared in ultrapure water or ethanol onto 300 mesh gold grids (Ted Pella) coated with lacey carbon and ultrathin carbon support films. Prior to deposition, the grids were rendered hydrophilic by glow-discharge treatment (25 mA, 50 s) using a Pelco easiGlow™ system. After deposition, the grids were dried under vacuum (∼ −250 mmHg). Particle size, morphology, crystallinity, and lattice fringes were evaluated from TEM and HRTEM images using ImageJ. For particle size analysis, four representative images from different regions of the sample were analyzed to obtain more reliable and representative measurements. Elemental composition and spatial distribution of chemical elements were investigated by Energy-Dispersive X-ray Spectroscopy (EDS) coupled to the STEM detector.

### 2.4 Biological Assays

#### 2.4.1 Cell Culture

The American Type Culture Collection (ATCC)-derived MDA-MB-231 (HTB-26) and SKBR3 (HTB-30) cell lines were cultured in complete DMEM. The medium was supplemented with 10% (v/v) fetal bovine serum (FBS), 4 mM glutamine, and penicillin (100 U/mL)/streptomycin (100 µg/mL). MCF-7 (HTB-22) cells were maintained in complete RPMI-1640 medium (Sigma-Aldrich) supplemented with the same components, except for glutamine, which was added at a final concentration of 2 mM. All cell lines were maintained under normoxic conditions (21% O₂) at 37 °C in a humidified atmosphere with 5% CO₂.

#### 2.4.2 Relative Metabolic Activity (MTT)

Cell viability was inferred by relative metabolic activity (% of control) sessed using the CyQUANT™ MTT kit (Invitrogen, Cat. V13154) according to the manufacturer’s instructions. SKBR3, MCF-7, and MDA-MB-231 cells were seeded in 96-well plates at a density of 1 × 10⁴ cells/well and incubated for 24 h at 37 °C under normoxic conditions. Cells were subsequently exposed to CuNP-PVP at concentrations ranging from 0.001 to 100 µg/mL for 48 h. After treatment, the culture medium was replaced with MTT solution, and the plates were incubated for 2.5 h. Formazan crystals were dissolved in DMSO, and absorbance was measured at 540 nm using a PerkinElmer EnSpire microplate reader (PerkinElmer).

Relative metabolic activity was calculated by normalizing the absorbance of treated samples to that of untreated control cells after background subtraction, according to the following equation:

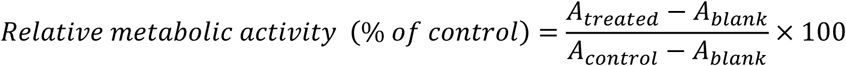

where A_treated_ corresponds to the absorbance of CuNP-treated cells, A_control_ represents the mean absorbance of untreated control cells, and A_blank_ corresponds to the absorbance of cell-free wells used for background correction.

The final analysis was performed using three independent biological replicates, each consisting of three to four technical replicates. For each cell line and CuNP concentration, cell viability was calculated as the mean of the corresponding technical replicates within each biological replicate. Data are presented as mean ± standard error of the mean (SEM), where bar heights represent the mean of biological replicate means and error bars indicate the SEM. Individual points correspond to the mean value of each biological replicate, allowing visualization of variability among biological replicates within each experimental group.

#### 2.4.3 MTT’s Statistical Analysis

Statistical analyses were performed using the non-parametric Kruskal–Wallis test to compare cell viability among the three breast cancer cell lines (MDA-MB-231, SKBR3, and MCF-7) at each CuNP concentration. When a significant overall effect was detected, pairwise comparisons were conducted using the Conover–Iman post-hoc test with Holm–Bonferroni correction for multiple testing. Differences were considered statistically significant at p < 0.05.

#### 2.4.4 Transcriptomic Analysis and Construction of Latent Copper-Response Programs

RNA-seq data from the luminal breast cancer cell line MCF-7 and the basal-like/triple-negative breast cancer cell line MDA-MB-231 were obtained from the GSE75168 dataset [42]. Raw HTSeq count matrices were analyzed using DESeq2 in R (v4.5.1). Count data were normalized using variance stabilizing transformation (VST) and the resulting expression matrix was used for principal component analysis (PCA), differential expression analysis, and heatmap generation. For Reactome and Hallmark pathway enrichment analyses, Gene Set Enrichment Analysis (GSEA) was performed using raw data as input.

Three latent copper-response gene programs were defined using genes associated with mitochondrial lipoylation, antioxidant defense, and lysosomal adaptation. The Lipoylation program included *LIAS*, *LIPT1*, *GCSH*, and *DLST*. The Antioxidant Defense program included *NFE2L2*, *KEAP1*, *GCLM*, *HMOX1*, *CAT*, *TXN*, *TXNRD1*, and *GPX4*. The Lysosomal Adaptation program included *TFEB*, *MCOLN1*, *CTSB*, *RAB27A*, *LAMP1*, and *LAMP2*. Heatmaps were generated using VST-normalized expression values subjected to gene-wise z-score normalization. Program scores were calculated as the arithmetic mean expression of all genes included in each signature.

Transcriptomic data from breast cancer cell lines were obtained from the Cancer Cell Line Encyclopedia (CCLE) through DepMap. Expression values corresponded to log₂ (TPM + 1). Cell lines were classified as Luminal, HER2-positive, or Basal/TNBC according to molecular subtype annotations. Lipoylation, Antioxidant Defense, and Lysosomal Adaptation scores were calculated using the same gene signatures defined in the discovery dataset. Subtype-level heatmaps were generated using row-wise z-score-standardized average scores.

Transcriptomic and clinical data from primary breast tumors were obtained from the TCGA-BRCA cohort through UCSC Xena. Gene-expression values were retrieved from the TCGA. BRCA,sampleMap/HiSeqV2 dataset, and PAM50 subtype classifications were obtained from the corresponding clinical annotations. Tumors were classified as Luminal A (LumA), Luminal B (LumB), HER2-enriched (Her2), or Basal-like (Basal). Program scores were calculated using the same gene signatures and scoring procedure applied to the cell-line datasets.

Potential Biological Copper-Response Index (PBCRI) was calculated by integrating the three transcriptional programs. Program scores were standardized across samples using z-score normalization, and the PBCRI was defined as:

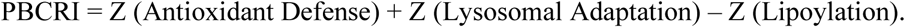

Positive PBCRI values indicate enrichment of antioxidant and lysosomal programs relative to lipoylation, whereas negative values indicate relative enrichment of the lipoylation program.

Comparisons among molecular subtypes in the CCLE and TCGA cohorts were performed using two-sided Wilcoxon rank-sum tests with Benjamini–Hochberg correction for multiple testing. All analyses were conducted in R using the packages DESeq2, dplyr, tidyr, ggplot2, pheatmap, RColorBrewer, UCSCXenaTools, and data.table.

## 3. Results and Discussion

### 3.1. Synthesis and Characterization of Polyvinylpyrrolidone-Copper Nanoparticles

#### 3.1.1. Formation, Optical and Colloidal Characterization of CuNP-PVP

To investigate the physicochemical properties of the synthesized CuNP-PVP, a comprehensive characterization was performed using spectroscopic, microscopic, and surface-sensitive techniques. UV–Vis spectroscopy showed that a localized surface plasmon resonance (LSPR), characteristic of metallic copper nanoparticles, emerged only after the two-step reduction process. After NaBH₄ reduction, no LSPR band was detected despite a visible color change (Figure 1A–I), indicating the formation of small copper nuclei incapable of supporting plasmonic oscillations. This is consistent with the rapid reduction kinetics of NaBH₄, which favor high nucleation density while limiting particle growth [43].

**Figure 1.**
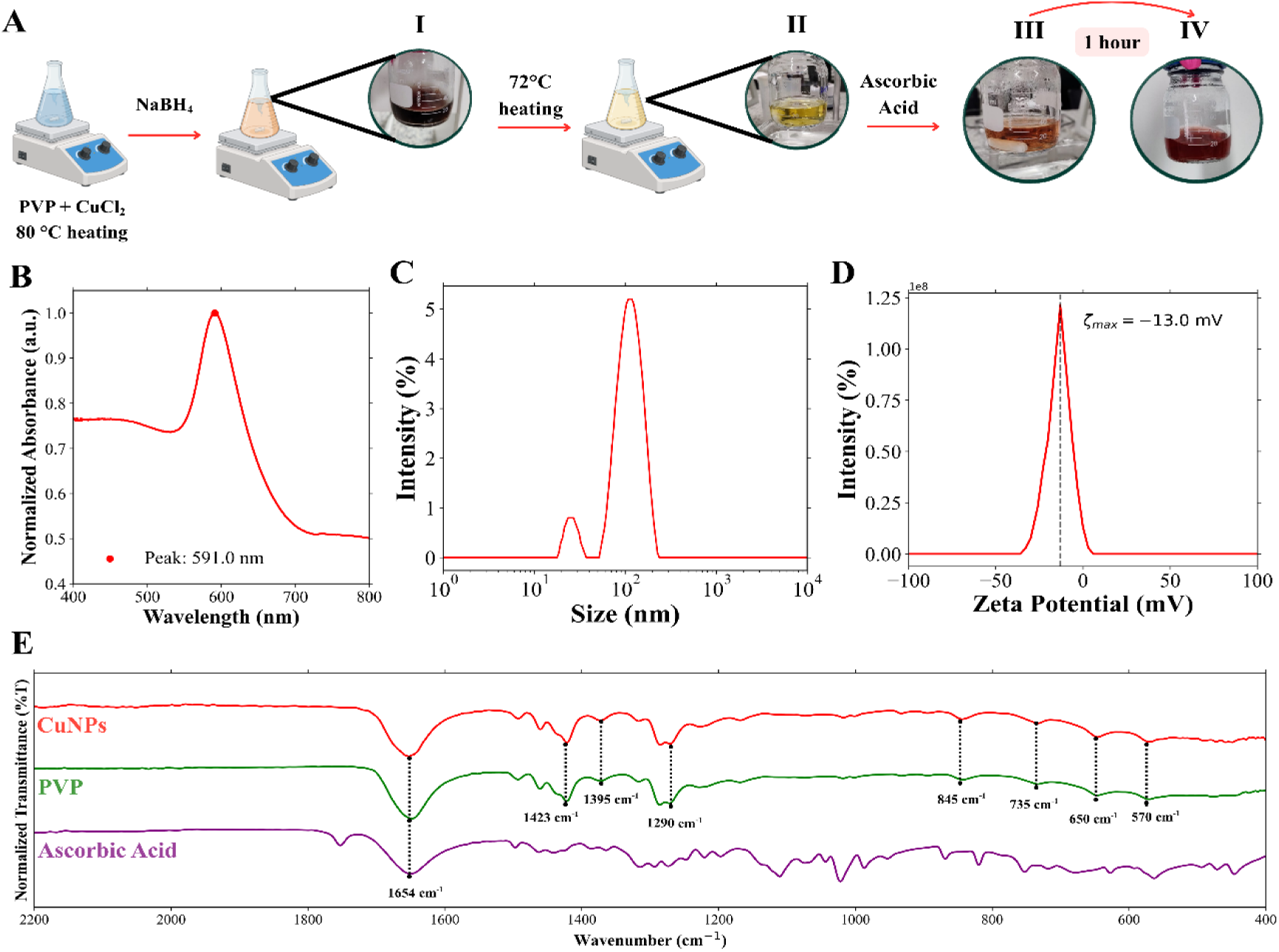
Synthesis and physicochemical characterization of PVP-assisted copper nanoparticles (CuNP-PVP). (A) Schematic representation of the synthesis process, showing the reaction stages: (I) immediately after NaBH₄ addition, (II) after resuming heating following ice-bath cooling, (III) after ascorbic acid addition, and (IV) the final nanoparticle suspension obtained after 1 h at room temperature. (B) UV–Vis absorption spectrum displaying a plasmon resonance band centered at 591 nm. (C) DLS size distribution, indicating nanoparticles with a bimodal distribution profile near 10 nm and centered at 100 nm. (D) Zeta potential distribution with a maximum at −13.0 mV. (E) FTIR spectra of CuNP-PVP, PVP, and ascorbic acid, confirming the presence of surface-bound stabilizing molecules on the nanoparticles.

An intermediate oxidized stage was observed before ascorbic acid addition (Figure 1A-II), suggesting partial oxidation of the initially formed nuclei [44]. Upon addition of the second reducing agent, the suspension underwent an abrupt color transition (Figure 1A-III), followed by the development of the characteristic reddish coloration characteristic of metallic copper colloids (Figure 1A-IV) and the appearance of a pronounced LSPR band centered at approximately 591 nm (Figure 1B). The emergence of this plasmonic feature confirmed the formation of metallic Cu domains within the colloid, while its broad and slightly red-shifted profile suggested size heterogeneity, interparticle interactions, and partial surface oxidation within the colloidal population [45].

To investigate the origin of this broadened plasmonic response, we analyzed particle size distribution by DLS (Figure 1C). The analysis revealed a bimodal profile comprising a nanoparticle population near 10 nm and a larger agglomerated population centered around 100 nm. This coexistence of nanoparticles and aggregates likely contributes to the optical heterogeneity observed by UV–Vis spectroscopy. Despite aggregation, the low polydispersity index (PDI < 0.3) indicated moderate uniformity within the dispersed populations. We next evaluated whether the purified nanoparticles retained a stable surface coating. ζ-potential measurements showed a moderately negative surface charge (≈ −13 mV; Figure 1D), likely reflecting partial removal of excess PVP during purification and consequent exposure of the underlying, possibly oxidized, nanoparticle surface. The low absolute zeta potential may be attributed to the combined effects of PVP coating and the adsorption of ionic species from the surrounding medium, such as OH⁻ and other negatively charged species potentially remaining from the synthesis. As a nonionic polymer, PVP can partially shield the intrinsic surface charge of the nanoparticles, while also providing steric stabilization that contributes to colloidal stability [46].

Consistently, FTIR analysis confirmed the presence of PVP after purification (Figure 1E). The spectrum exhibited the characteristic vibrational bands of the polymer, including the carbonyl stretching of the pyrrolidone ring (∼1650 cm⁻¹) and C–N/CH₂ vibrations between 1450 and 1280 cm⁻¹ [35]. No characteristic bands attributable to residual ascorbic acid were detected after purification, suggesting effective removal of unbound reagent.

Together, UV–Vis and DLS analyses indicated the formation of CuNP-PVP with heterogeneous size distribution and partial aggregation, whereas ζ-potential and FTIR measurements supported the persistence of the PVP coating after purification. Although these techniques provided important information regarding nanoparticle formation, colloidal properties, and surface functionalization, complementary analyses were realized to directly evaluate particle morphology, crystallinity, oxidation state, and elemental distribution.

#### 3.1.2 Morphological and Compositional Analysis

STEM imaging revealed predominantly spherical CuNP-PVP organized in agglomerated assemblies on the carbon support film (Figure 2A, Figure S1). Despite cluster formation, individual particle boundaries remained well defined, indicating limited particle coalescence during synthesis. Particle size analysis yielded an average diameter of 76.1 ± 21.0 nm (n = 179) (Figure 2B), consistent with the nanoscale population detected by DLS and supporting the broad optical response observed by UV–Vis spectroscopy. Furthermore, the larger hydrodynamic population observed is likely associated with nanoparticles coated by macromolecules [47,48] and could be related to agglomerations in the system.

**Figure 2.**
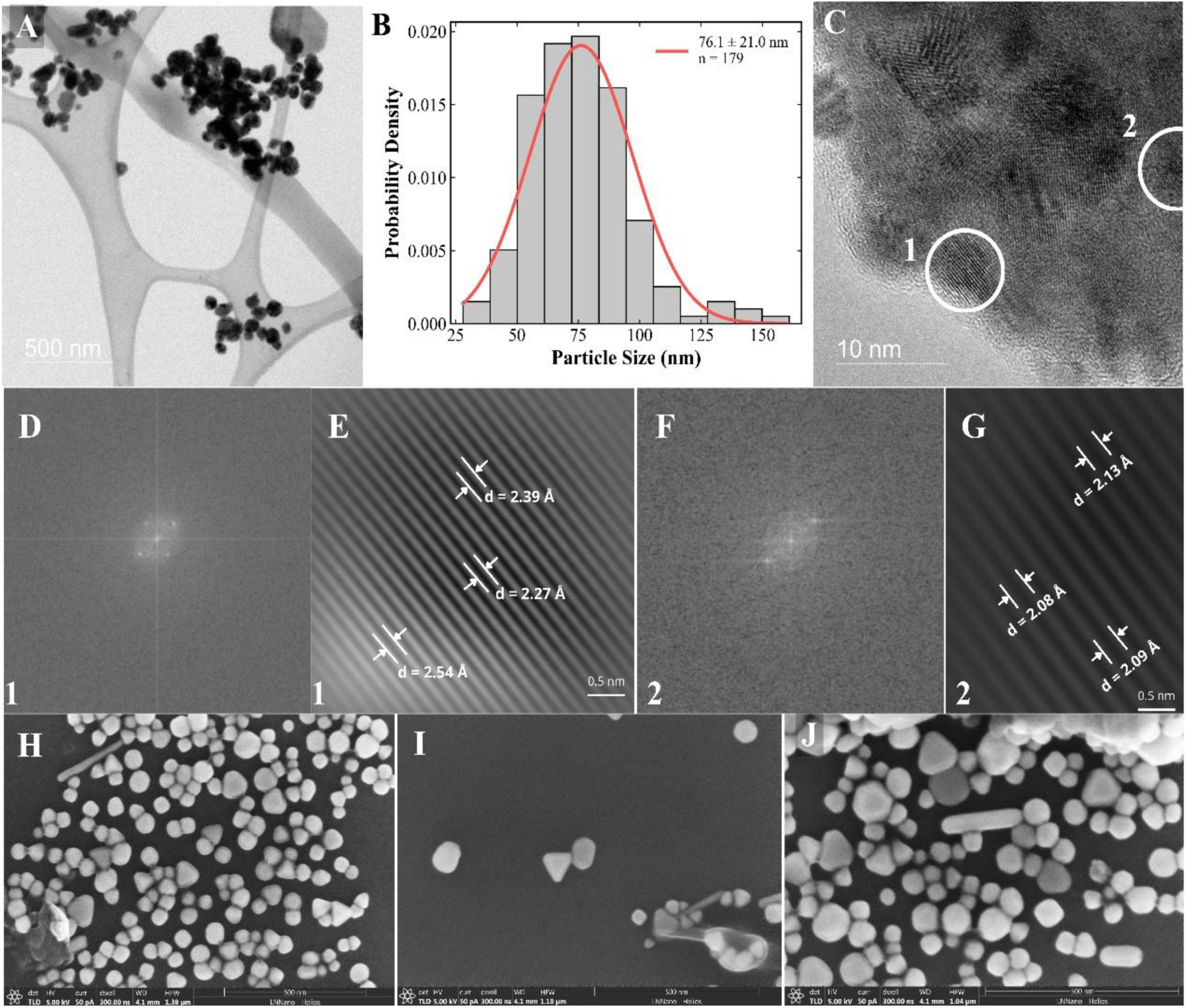
Morphological and structural characterization of CuNP–PVP. (A) Low-magnification STEM image showing predominantly spherical nanoparticles dispersed on the carbon support film, with localized agglomeration. (B) Particle size distribution determined from STEM images (n = 179), yielding an average diameter of 76.1 ± 21.0 nm. (C) HRTEM image highlighting two representative crystalline domains selected for lattice analysis. (D, F) Fast Fourier transform (FFT) patterns obtained from domains 1 and 2, respectively, confirming their crystalline nature. (E, G) Corresponding inverse FFT images revealing well-defined lattice fringes with interplanar spacings of 2.54, 2.39, and 2.27 Å in domain 1, and 2.13, 2.09, and 2.08 Å in domain 2. The spacing of ∼2.09 Å is consistent with the Cu(111) plane of face-centered cubic metallic copper, whereas the larger spacings are attributed to copper oxide phases (Cu₂O/CuO), indicating the coexistence of metallic and oxidized copper species. (H-J) SEM images acquired from different regions of the sample, showing predominantly spherical particles together with a smaller population of faceted and anisotropic morphologies.

HRTEM images revealed well-defined lattice fringes, confirming the crystalline nature of the CuNP-PVP samples (Figure 2C). Fast Fourier transform (FFT) patterns obtained from regions 1 and 2 are shown in Figs. 2D and 2F, respectively. Inverse FFT analysis of region 1 (Figure 2E), yielded interplanar spacings of 2.54, 2.39, and 2.27 Å. The 2.54 Å spacing is consistent with the CuO {111}/{−111} reflections (ICDD PDF 48-1548), whereas the 2.39 Å spacing is compatible with the Cu₂O {111} plane (ICDD PDF 05-0667). The 2.27 Å spacing could not be unambiguously assigned based on lattice spacing alone. Similarly, inverse FFT analysis of region 2 (Figure 2G), revealed lattice spacings of 2.13 and 2.09 Å, corresponding to the Cu₂O {200} (ICDD PDF 05-0667) and Cu {111} planes (ICDD PDF 04-0836), respectively. Although definitive phase assignment would require complementary diffraction analyses, the measured lattice spacings strongly suggest the coexistence of metallic copper and copper oxide phases in the same particle.

The structural heterogeneity identified by HRTEM was also reflected in the particle morphology observed by SEM (Figs. 2H–J). While spherical nanoparticles predominated, a smaller fraction of faceted and anisotropic structures were detected. This morphological diversity is consistent with the broad size distribution observed by STEM and DLS and may reflect variations in growth kinetics during the two-step reduction process as well as facet-dependent interactions between PVP and the nanoparticle surface [49]. Together, the electron microscopy analyses revealed crystalline CuNP-PVP with heterogeneous morphologies and mixed metallic/oxidized domains. To further investigate the chemical composition and surface properties of these nanoparticles, complementary XPS and EDS analyses were performed.

#### 3.1.3 Surface Chemical Characterization

To further investigate the surface composition of CuNP-PVP, XPS analysis was performed (Figure 3). The Cu 2p spectrum (Figure 3A) exhibited the characteristic Cu 2p₃/₂–Cu 2p₁/₂ doublet [50]. The main Cu 2p₃/₂ peak centered at 932.6 eV is characteristic of reduced copper species (Cu⁰/Cu⁺), whereas the shake-up satellites observed between 940–945 eV and 960–965 eV indicate the presence of Cu²⁺ species [51]. These results support the coexistence of reduced and oxidized copper species on the nanoparticle surface, in agreement with the HRTEM observations.

**Figure 3.**
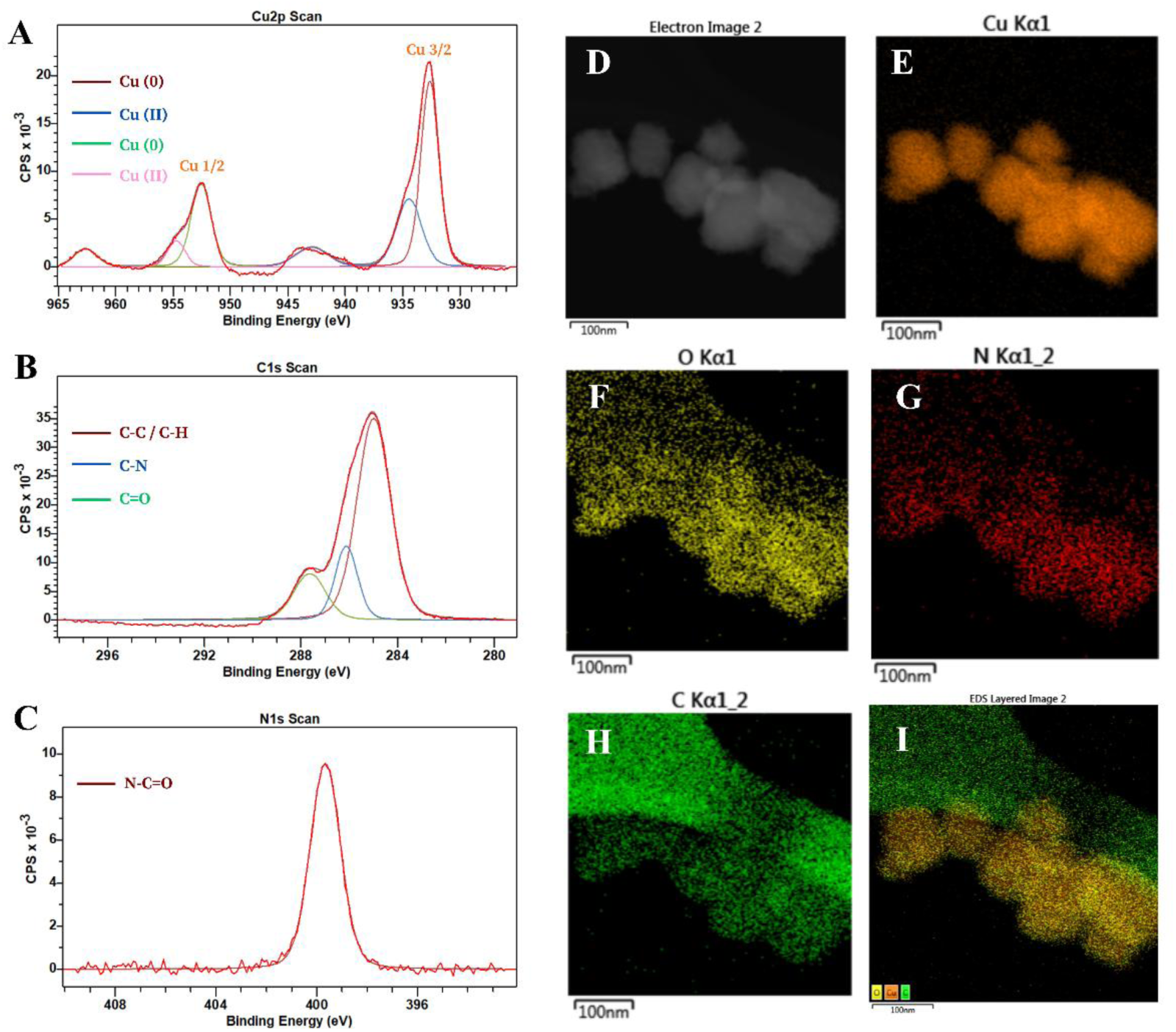
XPS and EDS characterization of CuNP-PVP. (A) Cu 2p XPS spectrum revealing the coexistence of reduced (Cu⁰/Cu⁺) and oxidized (Cu²⁺) copper species. (B,C) C 1s and N 1s spectra confirming the presence of PVP through characteristic carbon and amide nitrogen environments. (D) STEM image used for elemental mapping analysis. (E–H) EDS maps showing the distribution of Cu, O, N, and C. Copper is homogeneously distributed throughout the aggregates, whereas the overlap of O, N, and C signals supports the association of PVP with nanoparticles containing partially oxidized surface domains. (I) Composite elemental map highlighting the co-localization of Cu, O, and N signals.

The presence of PVP on the nanoparticle surface was further evaluated through analysis of the C 1s (Figure 3B) and N 1s (Figure 3C) spectra. Deconvolution of the C 1s region revealed contributions assigned to C–C/C–H, C–N, and carbonyl (C=O) environments, which are characteristic of the PVP molecular structure [52]. In addition, the N 1s spectrum displayed a component centered near 400 eV, consistent with the amide nitrogen present in the pyrrolidone ring. The O 1s spectrum (Supplementary Figure 1) exhibited contributions in the 531–532 eV region; however, because oxygen may originate from both copper oxide species and the carbonyl groups of PVP, a definitive assignment of this signal to a specific chemical environment is not straightforward.

While XPS provided information regarding copper oxidation states and surface chemistry, it does not reveal elemental distribution. Therefore, EDS elemental mapping was performed (Figs. 3E–I). Copper was uniformly distributed throughout the nanoparticle aggregates, whereas oxygen and nitrogen signals spatially overlapped with copper-rich regions. Together with the layered elemental map (Figure 3I), these observations support the association of the PVP coating with nanoparticles containing partially oxidized copper domains. Collectively, the physicochemical characterization confirmed the successful synthesis of CuNP-PVP with a predominantly spherical morphology, nanoscale dimensions, and the coexistence of reduced and oxidized copper species. Given the known influence of nanoparticle physicochemical properties on biological activity, we next evaluated the effects of CuNP-PVP in breast cancer cell lines representing distinct molecular subtypes.

### 3.2 Biological Evaluation

#### 3.2.1 Cell-line dependent MTT Responses to CuNP-PVP

Exposure to CuNP-PVP produced distinct relative metabolic activity profiles across breast cancer cell lines (Figure 4A). Viability remained largely unaffected at concentrations between 0.001 and 10 μg/mL. Significant subtype-specific differences emerged only at 100 μg/mL (Kruskal–Wallis, p = 0.03; Figure 4B). At this concentration, MCF-7 cells exhibited the lowest viability, whereas MDA-MB-231 and SKBR3 cells showed higher survival rates. Pairwise comparisons confirmed significantly higher viability in MDA-MB-231 and SKBR3 cells than in MCF-7 (Figure 4C). No significant differences were detected between MDA-MB-231 and SKBR3.

**Figure 4.**
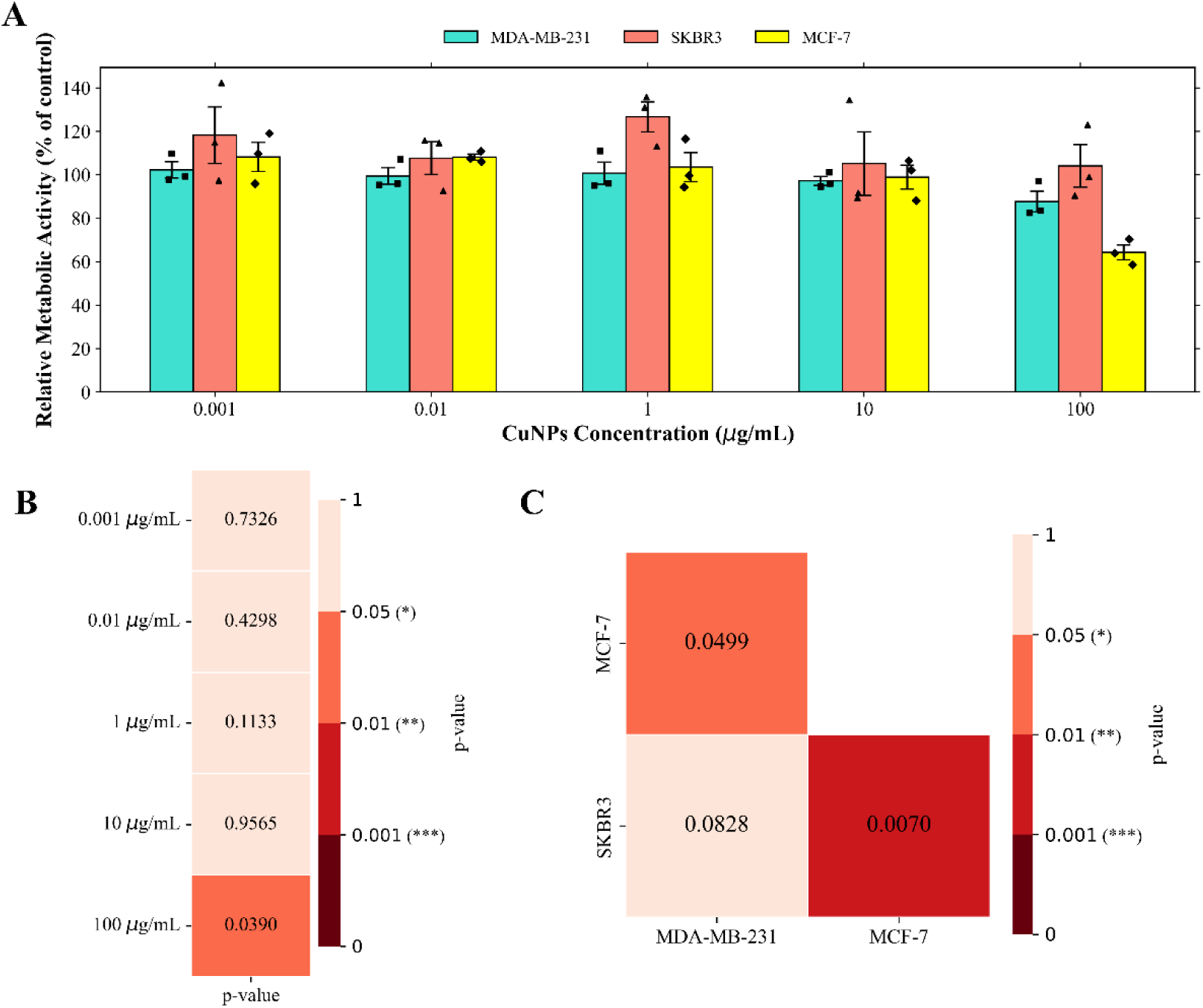
Differential sensitivity of breast cancer cell lines to CuNP-PVP exposure. (A) Cell viability was inferred by relative metabolic activity (% of control) measured by MTT assay following treatment with increasing concentrations of CuNP-PVP. Cell-specific differences became evident at 100 μg/mL, with MCF-7 exhibiting lower viability than MDA-MB-231 and SKBR3 cells. Bars represent the mean ± SEM of independent biological replicates, and symbols indicate individual biological replicate means. (B) Heatmap of Kruskal–Wallis p-values comparing viability across cell lines at each concentration, showing a significant effect only at 100 μg/mL. (C) Heatmap of pairwise Conover–Iman post hoc comparisons with Bonferroni correction for the 100 μg/mL treatment condition. Significant differences were observed between the more sensitive (MCF-7) and more tolerant (MDA-MB-231 and SKBR3) cell lines.

These findings are consistent with previous reports showing that luminal breast cancer cells are generally more susceptible to copper-induced cell death than TNBC models following treatment with copper ionophores [50]. Because the concentration range evaluated in the present study did not reach 50% inhibition, an IC₅₀ could not be reliably estimated. Nevertheless, the concentration-dependent reduction in MCF-7 viability agrees with the trend reported by Sharma et al. [41], who evaluated PVP-CuNPs using the same concentration range (0.001–100 μg/mL). Although their nanoparticles produced a stronger cytotoxic effect at the highest concentrations, both studies showed minimal effects at low doses and a dose-dependent increase in cytotoxicity. Future studies could expand the concentration range and time points and investigate IC_50_ for subtype-dependent sensitivity.

#### 3.2.2 Baseline Transcriptomic Differences Between Luminal and Triple-Negative Breast Cancer Models

To explore transcriptional features relevant to cell line-dependent MTT Responses to CuNP-PVP, we analyzed public baseline RNA-seq data (GSE75168) from MCF-7 and MDA-MB-231 cells. These lines exhibited contrasting sensitivity profiles and represented well-established luminal and TNBC models, respectively. Principal component analysis (PCA) showed tight clustering of biological replicates and clear separation between cell lines (Figure 5A). Differential expression analysis further revealed extensive transcriptional divergence between the two models (Figure 5B).

**Figure 5.**
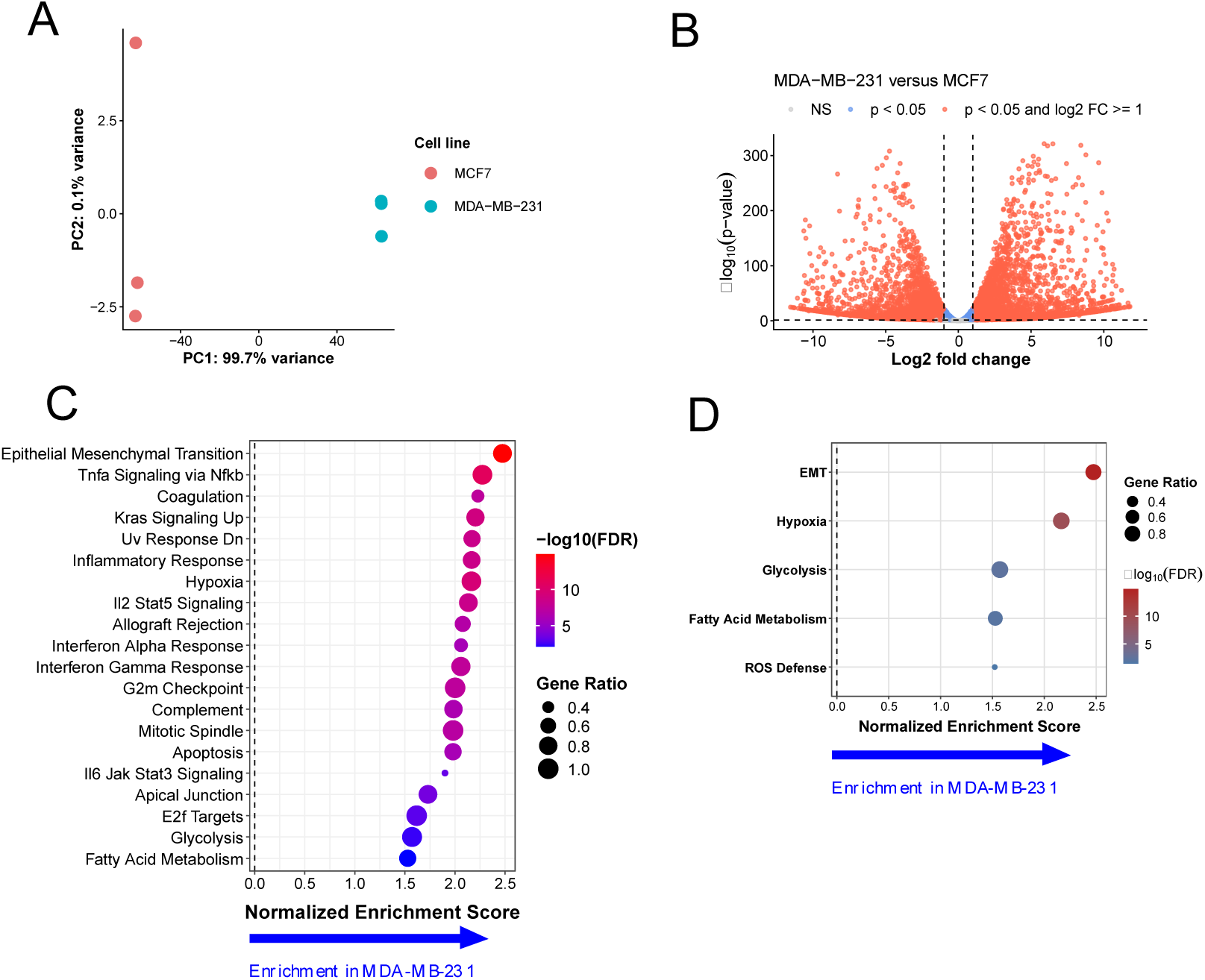
Transcriptomic programs possibly associated with differential copper nanoparticle response in breast cancer cells. (A) Principal component analysis (PCA) of transcriptomic profiles from MCF7 and MDA-MB-231 breast cancer cells, demonstrating segregation between luminal and basal-like phenotypes. (B) Volcano plot showing differentially expressed genes between MDA-MB-231 and MCF7 cells. (C) Hallmark gene set enrichment analysis (GSEA) of differentially expressed genes. (D) Hallmark pathways possibly associated with copper response identified by GSEA, showing normalized enrichment scores (NES), false discovery rates (FDR), and gene ratios for significantly enriched biological processes.

To further characterize the biological differences between the two cell lines, we performed Gene Set Enrichment Analysis (GSEA) using the Hallmark and Reactome gene set libraries. Hallmark analysis revealed enrichment of epithelial–mesenchymal transition (EMT), inflammatory, proliferative, and stress-associated programs in MDA-MB-231 cells, including TNFα/NFκB signaling, hypoxia, KRAS signaling, and cell-cycle-related pathways (Figure 5C).

Given our interest in the differential response to CuNP-PVP, we next examined pathways potentially relevant to copper-induced stress and cellular adaptation. A selected view of the Hallmark results highlighted enrichment of ROS response, hypoxia, glycolysis, fatty acid metabolism, and EMT programs in MDA-MB-231 cells (Figure 5D). The TNBC-enriched pathways are strongly associated with adaptation to metabolic and oxidative stress [54,55]. Enhanced ROS defense may improve TNBC cells’ ability to cope with copper-induced oxidative damage, while enrichment of hypoxia, glycolysis, and fatty acid metabolism suggests greater metabolic flexibility [56,57]. Likewise, EMT has been linked to cellular plasticity and increased tolerance to environmental stressors [58].

Reactome analysis further revealed enrichment of extracellular matrix organization, cell adhesion, cell motility, and mechanotransduction in MDA-MB-231 cells (Supplementary Figure 2A). These pathways are commonly associated with cellular adaptation, phenotypic plasticity, and environmental stress responses [59]. Collectively, the GSEA results highlight differences in metabolic, stress-associated, and cell-interaction programs between MDA-MB-231 and MCF-7 cells, providing a basis for investigating how these features may influence their responses to CuNP-PVP.

#### 3.2.3 Candidate Copper-Response Programs Identified by Transcriptomic Analysis

The enrichment analyses described above suggested that MDA-MB-231 cells possess a basal transcriptional landscape associated with metabolic and oxidative stress responses. Based on these results, we next examined three biological programs potentially relevant to copper responses: protein lipoylation, antioxidant defense, and lysosomal adaptation.

Analysis of representative genes revealed distinct expression patterns between the two cell lines (Figure 6A). Genes involved in protein lipoylation, including *LIAS*, *LIPT1*, *GCSH*, and *DLST*, were more highly expressed in MCF-7 cells than in MDA-MB-231. This pattern is consistent with current models of cuproptosis, which depend on lipoylated mitochondrial proteins [15,60]. In contrast, genes associated with antioxidant defense (*NFE2L2*, *KEAP1*, *GCLM*, *HMOX1*, *CAT*, *GPX4*, *TXN*, and *TXNRD1*) and lysosomal adaptation (*TFEB*, *MCOLN1*, *RAB27A*, *CTSB*, *LAMP1*, and *LAMP2*) were upregulated in MDA-MB-231 cells. To further illustrate the differences underlying these transcriptional programs, we examined normalized transcript abundance for the selected genes. As shown in Figure S2B, the differences between MCF-7 and MDA-MB-231 cells were maintained when assessed using normalized counts rather than row-wise Z-scores.

**Figure 6.**
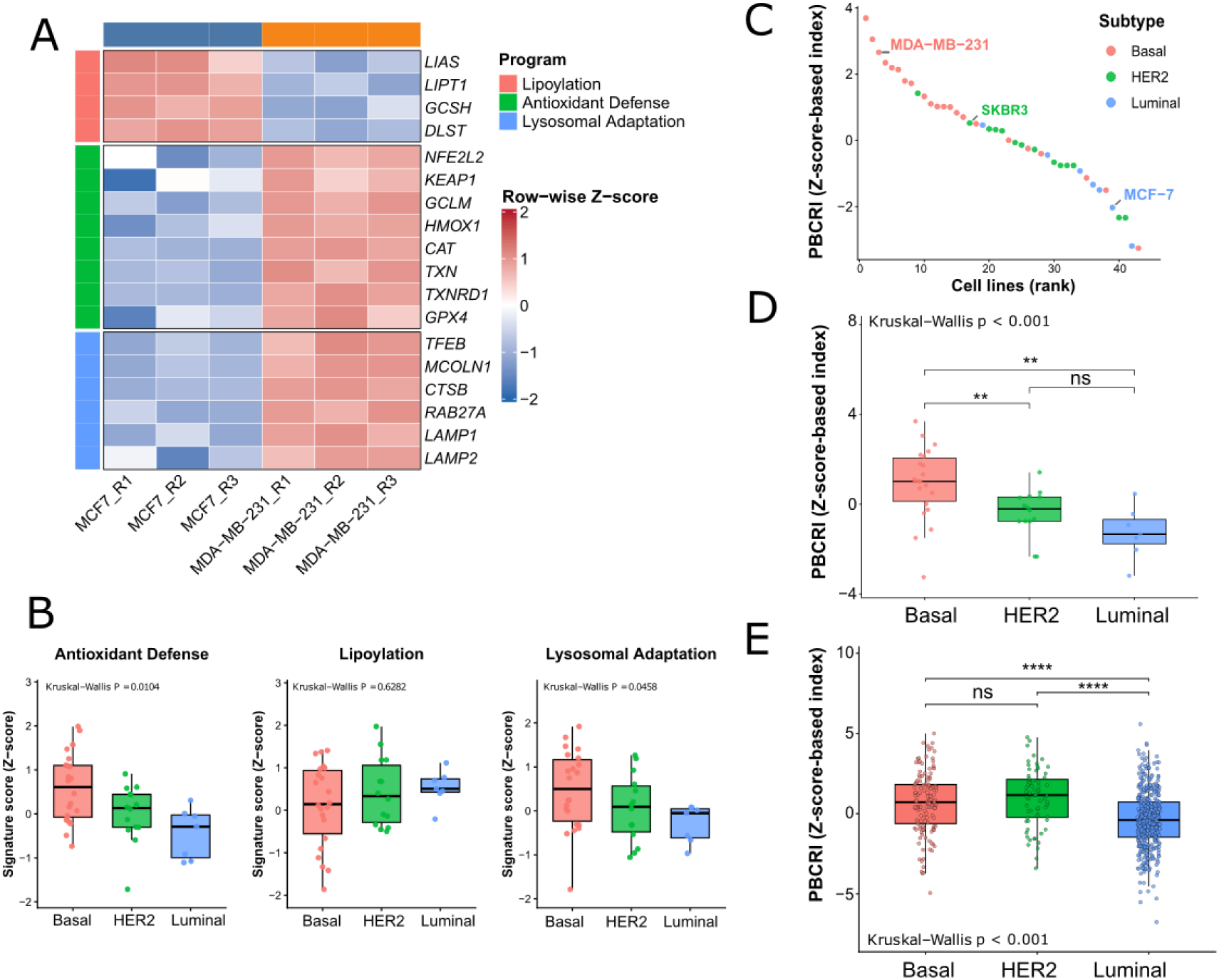
Latent Copper-response programs define adaptive states across breast cancer models and patient cohorts. (A) Heatmap showing the expression of genes associated with lipoylation, antioxidant defense, and lysosomal adaptation programs in MCF7 and MDA-MB-231 cells. (B) Activity scores of possibly copper-response programs across breast cancer molecular subtypes in the Cancer Cell Line Encyclopedia (CCLE). (C) Ranking of breast cancer cell lines according to the Potential Biological Copper-Response Index (PBCRI). Cell lines are ordered by PBCRI score and colored according to molecular subtype; representative MDA-MB-231, SKBR3, and MCF-7 models are indicated. (D) Potential Biological Copper Response Index (PBCRI) across breast cancer cell lines. The PBCRI was calculated as Z(AntioxidantScore) + Z(LysosomalScore) − Z(LipScore), where Z denotes the standardized score of each mechanistic program. Higher values indicate a more copper-adapted transcriptional state. (E) PBCRI across TCGA-BRCA molecular subtypes. Higher PBCRI values indicate increased antioxidant and lysosomal activity combined with reduced lipoylation. (Statistical comparisons in panels C–E were performed using two-sided Wilcoxon rank-sum tests with Benjamini–Hochberg correction for multiple testing. Statistical significance is indicated as *p < 0.05, **p < 0.01, ***p < 0.001, and ****p< 0.0001

Together, these findings indicate distinct basal transcriptional profiles related to mitochondrial lipoylation, oxidative stress management, and lysosomal function between the two cell lines. Such differences may influence how the cells respond to copper-induced stress and may contribute to differences in their susceptibility to copper-mediated cytotoxicity. However, whether these transcriptional features directly confer protection against copper stress or alter cuproptosis susceptibility requires functional validation. We therefore examined whether these patterns were also observed at the pathway level across additional cell lines.

#### 3.2.4 Exploratory Evaluation Across Breast Cancer Models and Patient Tumors

To determine whether the transcriptional programs associated with differential CuNP-PVP sensitivity extended beyond the MCF-7 and MDA-MB-231 models, we performed an exploratory analysis of transcriptomic data from a broader panel of breast cancer cell lines from the Cancer Cell Line Encyclopedia (CCLE). Antioxidant defense and lysosomal adaptation showed significant subtype-associated differences across the evaluated cell lines (Figure 6B), whereas lipoylation showed a consistent trend accordingly with the MCF7 vs MDA-MB-231 results.

Because subtype-associated differences extended to multiple biological programs, we integrated the three gene programs into a single composite metric, the Potential Biological Copper-Response Index (PBCRI). The PBCRI summarizes the relative balance between the lipoylation-associated program and the antioxidant and lysosomal adaptation programs identified in the transcriptomic data. We then calculated the PBCRI for individual cell lines and ranked the models according to their signature scores. Basal-like cell lines were predominantly positioned toward the higher end of the PBCRI distribution, whereas luminal-like models tended to occupy the lower end, with HER2-like models generally distributed between the two groups (Figure 6C).

We next assessed whether PBCRI scores differed across molecular subtypes. A Kruskal–Wallis test indicated a significant overall difference among basal-like, HER2-like, and luminal-like cell lines (p < 0.001; Figure 6D). Basal-like models exhibited significantly higher PBCRI scores than both HER2-like and luminal-like models, whereas the difference between HER2-like and luminal-like models was not statistically significant. Collectively, these findings indicate that the transcriptional programs incorporated into the PBCRI are associated with breast cancer subtype, particularly distinguishing basal-like models from luminal-like and HER2-like models.

To assess whether these subtype-associated transcriptional patterns were also evident in clinical samples, we analyzed bulk RNA-seq data from TCGA-BRCA. Basal-like and HER2-enriched tumors showed higher antioxidant defense, lysosomal adaptation, and PBCRI scores than Luminal A tumors, whereas differences in the lipoylation program were less pronounced (Figure 6E; Figure Supplementary 2C). Notably, HER2-enriched tumors did not consistently exhibit an intermediate profile in the patient cohort, as their PBCRI distribution was comparable to or higher than that of basal-like tumors. Because TCGA-BRCA represents bulk tumor tissue, these scores may also reflect contributions from stromal and immune compartments.

Overall, the recurrence of subtype-associated transcriptional patterns across CCLE cell lines and TCGA-BRCA tumors indicates that the selected gene programs capture biological features associated with breast cancer subtype (Figure 6E). In particular, basal/TNBC models had higher antioxidant and lysosomal adaptation scores, whereas luminal models showed greater representation of the lipoylation program. These findings support the potential relevance of these transcriptional programs to basal differences in cellular responses to copper-induced stress. However, the PBCRI remains an exploratory composite signature derived from the MCF-7 versus MDA-MB-231 comparison, and its ability to predict copper or CuNP-PVP sensitivity will require independent functional validation.

## 4. Conclusion

In this exploratory study, we evaluated CuNP-PVP across breast cancer cell lines representing distinct molecular subtypes by integrating physicochemical characterization, cytotoxicity assessment, and basal transcriptomic analyses. The synthesized nanoparticles exhibited nanoscale dimensions, predominantly spherical morphology, mixed copper oxidation states, and a PVP coating. CuNP-PVP exposure induced cell-line-dependent effects at 100 μg/mL, with MCF-7 cells showing lower relative metabolic activity than SKBR3 and MDA-MB-231 cells. Baseline transcriptomic comparisons identified higher expression of selected lipoylation-associated genes in MCF-7 cells and higher antioxidant-defense- and lysosomal-associated scores in MDA-MB-231 cells. Similar subtype-associated score distributions were observed in CCLE and TCGA-BRCA datasets.

Integration of these transcriptional features into the PBCRI revealed subtype-associated patterns across independent breast cancer cell lines and patient tumors, with generally higher scores in basal-like models and lower scores in luminal models, while HER2-positive models showed a more variable profile. Together, these findings identify basal transcriptional features associated with breast cancer subtypes that may also be relevant to differential responses to copper-induced stress. The PBCRI provides an exploratory, hypothesis-generating framework for further investigating these associations, while its relationship with copper or CuNP-PVP sensitivity will require independent functional validation.

## Corresponding author

Cyro von Zuben de Valega Negrão., Brazilian Center for Research in Energy and Materials (CNPEM), Giuseppe Máximo Scolfaro Street, number 10.000, Polo II de Alta Tecnologia de Campinas, 13083-100, Campinas, Brazil.

## Acknowledgments

The authors acknowledge the Brazilian Center for Research in Energy and Materials (CNPEM) for providing the infrastructure that enabled this work, as well as the Ilum School of Science, where this project was initially developed, for its financial support and laboratory facilities. The authors also thank the Brazilian Biosciences National Laboratory (LNBio) for providing access to the research facilities where the biological assays were performed, and the Brazilian Nanotechnology National Laboratory (LNNano) for support with the microscopy and spectroscopy analyses. Finally, the authors thank all collaborators and colleagues who contributed to this project through valuable discussions, technical assistance, and ongoing support.

## Consent for publication

All authors consent to publication.

## Availability of data and materials

The datasets generated and/or analyzed during the current study are available from the corresponding author upon reasonable request.

## Author contribution

A. G. Silva, Izaque O. Silva, and C. Z. Valega Negrão contributed to conceptualization, experimental design, experimental execution, data analysis, manuscript drafting, and revision. P. P. Tanaka and C. Z. Valega Negrão contributed to transcriptomic analysis, manuscript drafting, and revision. M. Z. Monteiro contributed to experimental design, statistical analysis, manuscript drafting, and revision. S. S. O. Filho and J. F. M. Souza contributed to manuscript revision, while V. S. Marangoni contributed to experimental design and manuscript revision. S. Dias and C. Z. Valega Negrão supervised the study and critically revised the manuscript.

