## Supplementary_material for "Differential Cytotoxicity of PVP-Copper Nanoparticles in Breast Cancer Cell Lines: Insights from Basal Transcriptomic Profiles"

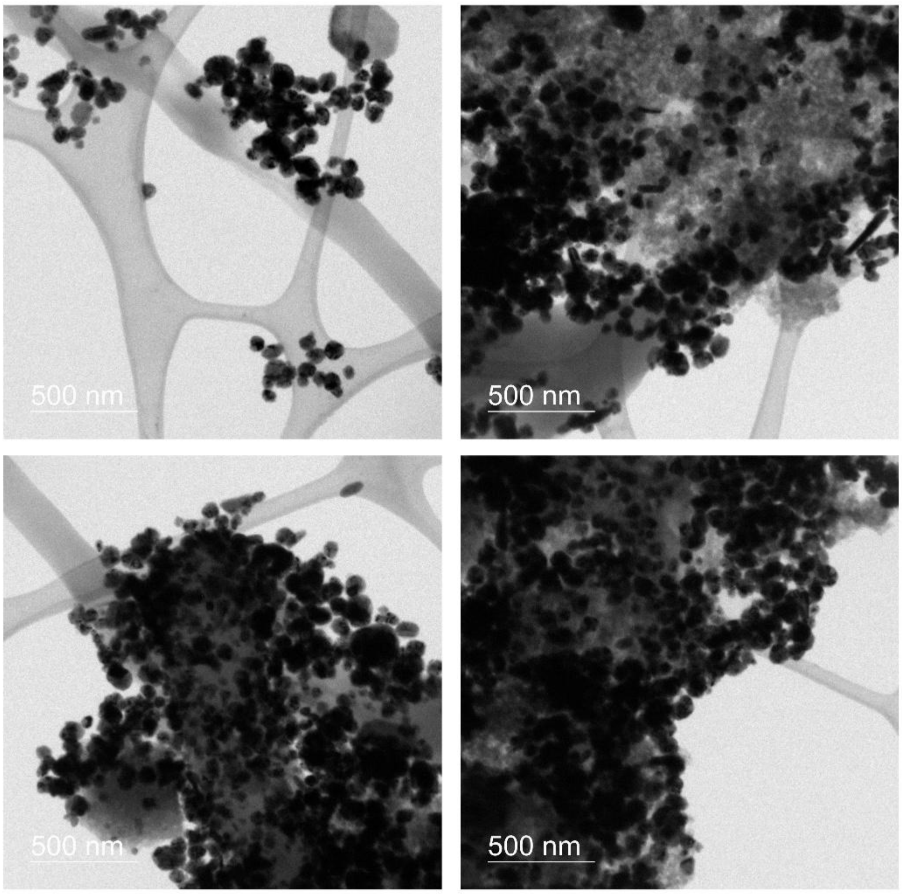


***Figure Supplementary 1.*** *Representative TEM micrographs of CuNP-PVP used for particle size analysis. The images were used to determine the size of individual nanoparticles using ImageJ, with the 500 nm scale bars used for image calibration.*

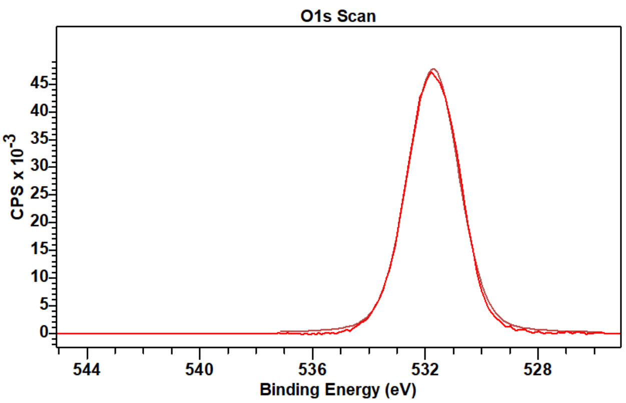


***Figure Supplementary 2.*** *X-ray photoelectron spectroscopy (XPS) spectrum of the O 1s region. The main peak centered at approximately 531.8 eV indicates the presence of oxygen-containing species on the sample surface. The black line represents the experimental data, while the red line corresponds to the fitted curve.*

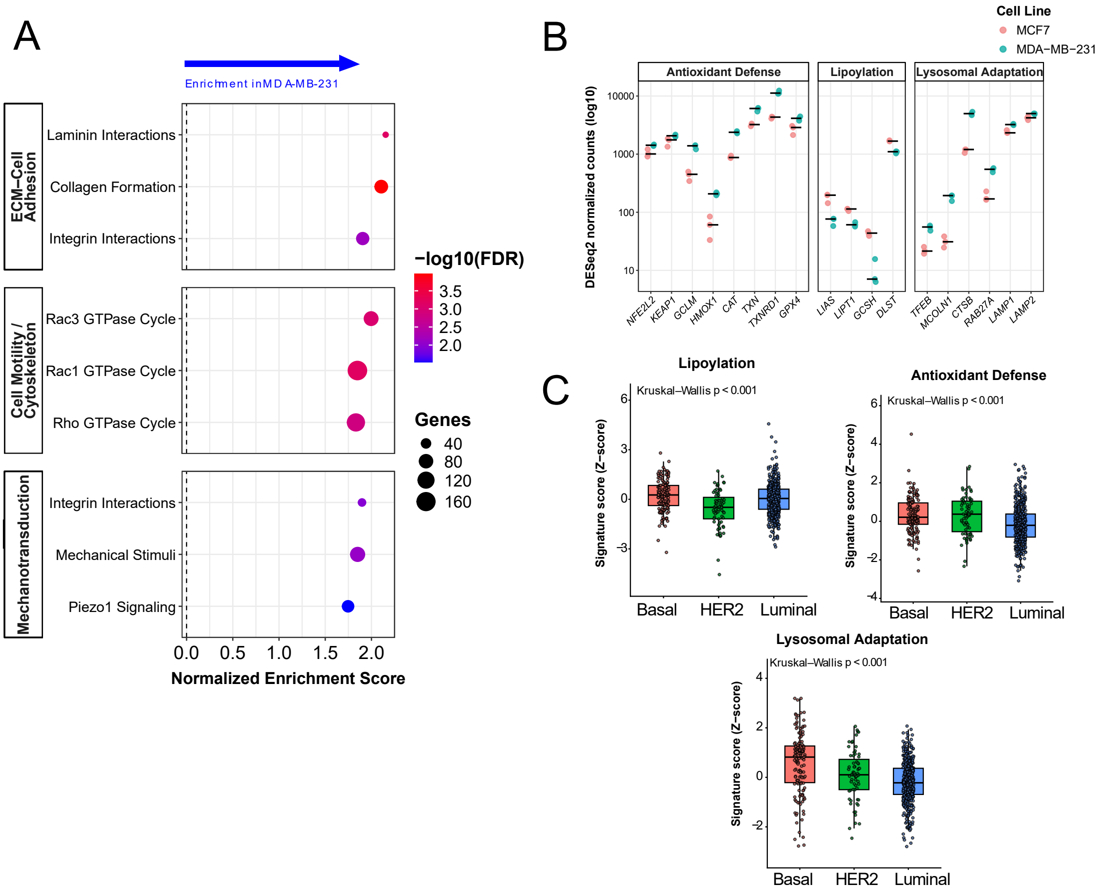


***Figure Supplementary 3.*** *(A) Reactome pathway enrichment analysis of genes differentially expressed between MCF7 and MDA-MB-231 cells. Significantly enriched pathways are shown by the normalized enrichment score and false discovery rate (FDR). (B) Expression of genes included in the Lipoylation (LIAS, LIPT1, GCSH, and DLST), Antioxidant Defense (NFE2L2, KEAP1, GCLM, HMOX1, CAT, TXN, TXNRD1, and GPX4), and Lysosomal Adaptation (TFEB, MCOLN1, CTSB, RAB27A, LAMP1, and LAMP2) programs across biological replicates of MCF7 and MDA-MB-231 cells. (C) Activity scores of Lipoylation, Antioxidant Defense, and Lysosomal Adaptation programs across TCGA-BRCA molecular subtypes. Statistical significance was assessed using two-sided Wilcoxon rank-sum tests with Benjamini–Hochberg correction. Statistical significance is indicated as *P < 0.05, **P < 0.01, ***P < 0.001, and ***P < 0.0001.*
